# Optogenetic activation of SST-positive interneurons restores hippocampal theta oscillation impairment induced by soluble amyloid beta oligomers *in vivo*

**DOI:** 10.1101/465112

**Authors:** Hyowon Chung, Kyerl Park, Hyun Jae Jang, Michael M Kohl, Jeehyun Kwag

## Abstract

Abnormal accumulation of amyloid β oligomers (AβO) is a hallmark of Alzheimer’s disease (AD), which leads to learning and memory deficits. Hippocampal theta oscillations that are critical in spatial navigation, learning and memory are impaired in AD. Since GABAergic interneurons, such as somatostatin-positive (SST+) and parvalbumin-positive (PV+) interneurons, are believed to play key roles in the hippocampal oscillogenesis, we asked whether AβO selectively impairs these SST+ and PV+ interneurons. To selectively manipulate SST+ or PV+ interneuron activity in mice with AβO pathology *in vivo*, we co-injected AβO and adeno-associated virus (AAV) for expressing floxed channelrhodopsin-2 (ChR2) into the hippocampus of SST-Cre or PV-Cre mice. Local field potential (LFP) recordings *in vivo* in these AβO–injected mice showed a reduction in the peak power of theta oscillations and desynchronization of spikes from CA1 pyramidal neurons relative to theta oscillations compared to those in control mice. Optogenetic-activation of SST+ but not PV+ interneurons in AβO–injected mice fully restored the peak power of theta oscillations and resynchronized the theta spike phases to a level observed in control mice. *In vitro* whole-cell voltage-clamp recordings in CA1 pyramidal neurons in hippocampal slices treated with AβO revealed that short-term plasticity of SST+ interneuron inhibitory inputs to CA1 pyramidal neurons at theta frequency were selectively disrupted while that of PV+ interneuron inputs were unaffected. Together, our results suggest that dysfunction in inputs from SST+ interneurons to CA1 pyramidal neurons may underlie the impairment of theta oscillations observed in AβO-injected mice *in vivo.* Our findings identify SST+ interneurons as a target for restoring theta-frequency oscillations in early AD.

## Introduction

Alzheimer’s disease (AD) is a neurodegenerative disease characterized by progressive memory loss and cognitive decline observed in humans as well as animal models of AD (Jahn, 2013; Kelley and Petersen, 2007; LaFerla and Green, 2012; LaFerla et al., 2007; Palop and Mucke, 2010, 2016). Although the causes of AD remain unknown, abnormal accumulation of soluble oligomers of amyloid β (AβOs) has been implicated in the pathogenesis of early stage of AD (Lambert et al., 1998; Lesne et al., 2006; McLean et al., 1999). AβOs have deleterious effects on neurons and synapses, causing synaptic dysfunction (Lacor et al., 2004; Shankar et al., 2008) and neural death (Alberdi et al., 2010; De Felice et al., 2007; Decker et al., 2010). Experimentally, AβO have been injected into the hippocampus *in vivo* to investigate the effect of AβO in hippocampal network oscillations, synaptic plasticity, and behavioral memory performance (Kalweit et al., 2015; Kim et al., 2014b; Nicole et al., 2016; Orban et al., 2010; Villette et al., 2010; Yi et al., 2018). In these studies, hippocampal theta oscillation, which is important in providing a clock mechanism for spike phase coding in spatial information processing (Dragoi and Buzsaki, 2006; Lisman and Jensen, 2013; O’Keefe and Recce, 1993) and support synaptic plasticity (Bikbaev and Manahan-Vaughan, 2008; Buzsaki, 2002; Huerta and Lisman, 1993; Nyhus and Curran, 2012; Tsanov and Manahan-Vaughan, 2009), was impaired in AD by showing decreased power compared to normal conditions. Distinct subtypes of hippocampal interneurons such as somatostatin-positive (SST+) (Fuhrmann et al., 2015) and parvalbumin-positive (PV+) interneurons are critically involved in the oscillogenesis (Wang and Buzsaki, 1996) and modulation of hippocampal theta oscillation (Amilhon et al., 2015; Huh et al., 2016). Recent experimental evidence showed that these interneurons disintegrate structurally and functionally in AD mice models (Palop et al., 2007; Schmid et al., 2016), suggesting that dysfunctions of SST+ and PV+ interneurons may underlie theta oscillation impairment observed in AD. Recent experiments attempted to find methods to restore impaired hippocampal gamma oscillation using magnetic stimulation (Zhen et al., 2017), by ablating ErbB4 in PV+ interneurons (Zhang et al., 2017), or by overexpressing Nav1.1 in interneurons (Martinez-Losa et al., 2018). However, no study has yet directly demonstrated whether modulation of SST+ and PV+ interneurons can restore impaired hippocampal theta oscillation in AD.

To address this, we selectively manipulated the activity in SST+ or PV+ interneurons using optogenetics in mice with AβO pathology *in vivo.* We co-injected AβO (Brouillette et al., 2012; Cetin and Dincer, 2007; Faucher et al., 2015; Kim et al., 2014b) and adeno-associated virus, AAV (AAV5-EF1a-DIO-hChR2-mCherry) for expressing floxed channelrhodopsin2 (ChR2) in SST+ and PV+ interneurons in SST-Cre mice or PV-Cre mice, respectively (Yizhar et al., 2011). Local field potential (LFP) recordings *in vivo* revealed that reduction of theta oscillation power and desynchronized spikes of CA1 pyramidal neurons relative to theta oscillation in AβO–injected mice were restored to the level observed in the control mice by optogenetic activation of SST+ interneurons with blue light but not by PV+ interneurons. *In vitro* whole-cell voltage-clamp recordings in CA1 pyramidal neurons in hippocampal slices treated with AβO revealed disruption of short-term plasticity of SST’s inhibitory inputs to pyramidal neurons at theta frequency. These results suggest a synaptic dysfunction of SST+ interneuron inputs to CA1 pyramidal neurons may underpin impairment of theta oscillation in AβO-injected mice *in vivo.*

## Methods

### Animals

SST-IRES-Cre (Jackson Laboratory, stock #013044, (Taniguchi et al., 2011)) knock-in mice and B6 PV-Cre (Jackson Laboratory, stock #017320) knock-in mice were housed in a temperature-controlled environment under a 12h light/dark cycle with the guideline of the Gyerim Experimental Resource Center of Korea University. Food and water were provided ad libitum. All experimental procedures were approved by the Institutional Animal Care and Use Committee (IACUC) of Korea University (KUIACUC-2017-112).

### Soluble AβO preparation

Soluble AβO was prepared following methods described by Lambert and colleagues (Lambert et al., 1998) with a slight modification (Wang et al., 2008). In brief, soluble Aβ_1-42_ was dissolved for monomerization in 1,1,1,3,3,3-hexafluoro-2-propanol (HFIP) at a final concentration of 1 mM and incubated for 90 min. HFIP was evaporated under vacuum condition in SpeedVac (N-BIOTEK Inc.). The remaining thin and clear film of Aβ_1-42_ was dissolved in dimethyl sulfoxide (DMSO) to make 5 mM Aβ_1-42_ stock, which was aliquoted and frozen at – 20 □. The Aβ_1-42_ stock was thawed and diluted to a final concentration of 200 nM in ariticial cerebrospinal fluid (aCSF). After dilution, Aβ_1-42_ solution was incubated for 18 hours for Aβ oligomerization.

### Stereotaxic surgery

In order to optogenetically manipulate SST+ or PV+ interneurons in mice with AβO pathology, we injected soluble AβO together with adeno-associated virus (AAV) vectors AAV5-EF1a-DIO-hChR2-mCherry (AAV5-EF1a-DIO-hChR2(E123T/T159C)-mCherry, 3.8×10^12^ virus molecules/mL, UNC Vector Core) in SST-Cre and PV-Cre mice (postnatal day 28 to 49) to express blue light-gated cation channel (Channelrhodopsin-2, ChR2) in PV+ and SST+ interneurons, respectively (Fig. 1A and Fig. 2A) (Fenno et al., 2011). SST-Cre and PV-Cre mice were deeply anesthetized with isoflurane and head-fixed into a stereotaxic frame (Stoelting Co.). The position of the hippocampal CA1 region (2.7 mm posterior, 2.7 mm lateral and 1.85 mm ventral to the bregma) was marked on the skull and thinned using a high-speed drill (K.1070, Foredom Electric Co.) to make a small craniotomy for injection. 3 μL of AβO_1-42_ (10 μM) (Kim et al., 2014a) and 1 μL of AAV5-EF1a-DIO-hChR2-mCherry were delivered into the hippocampal CA1 region through 5 μL micro-needle (Hamilton Company) connected to a motorized stereotaxic injector (Stoelting Co.) at the rate of 0.3 μL/min and 0.1 μL/min, respectively. Body temperature was maintained using a heating pad during surgery. After the injection, the needle was left in the brain for > 5 min to minimize backflow of the injection. At least three weeks of recovery period was allowed after surgery before *in vivo* recording to ensure the proper expression of the virus as well as induction of AβO–mediated oscillogenesis as observed in other study (Villette et al., 2010).

**Fig. 1.**
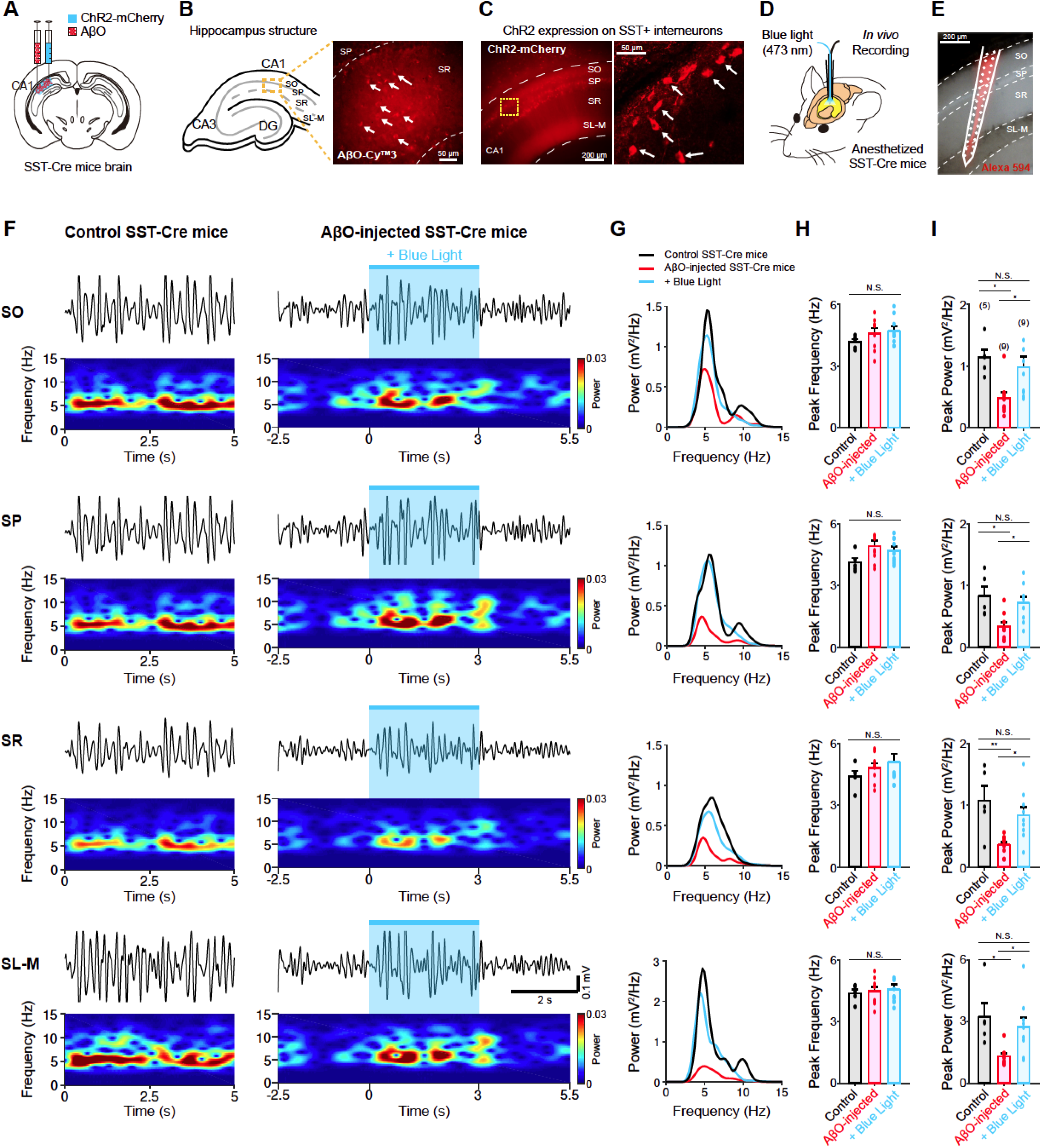
Optogenetic activation of SST+ interneurons in AβO-injected SST-Cre mice restored impaired hippocampal theta oscillation. (A) Schematic illustration of micro-injection of AβO and ChR2 virus into hippocampal CA1 region of SST-Cre mice. (B) Schematic illustration of hippocampal structure (left) and immunofluorescence image of AβO pointed by white arrows (right). (C) Fluorescence image of ChR2 expressed SST+ interneuron**s** in hippocampal CA1 region. Individual neurons are pointed by white arrows. (D) Schematic illustration of *in vivo* recording. (E) The recording location marked with Alexa 594 fluorescent dye and channel mapping of 32-channel silicon probe aligning with the Alexa 594 staining. The electrode crossed through stratum oriens (SO), stratum pyramidale (SP), stratum radiatum (SR) and stratum lacunosum-moleculare (SL-M) in CA1 area. (F) Example traces of spontaneous theta-filtered LFP signals (top) and spectrogram (bottom) recorded in control SST-Cre mice (left) and AβO-injected SST-Cre mice (right). The 3 s-long blue light stimulation of SST+ interneurons is represented by shaded region (light blue). (G) Example traces of power spectrum density (PSD) curves obtained from the spontaneous theta-filtered LFP signals recorded in control SST-Cre mice (black), AβO-injected SST-Cre mice (red) and AβO-injected SST-Cre mice with blue light (blue). (H-I) Mean peak frequency (H) and mean peak power (I) of spontaneous theta-filtered LFP signals of control SST-Cre mice (black, n = 5), AβO-injected SST-Cre mice (red, n = 9) and AβO-injected SST-Cre mice with blue light (blue, n = 9). All data represent mean ± SEM. Inset: N.S. *p* > 0.05, ^∗^ *p* < 0.05 and ^∗∗^ *p* < 0.01, one-way ANOVA followed by Tukey’s *post hoc* test.

**Fig. 2.**
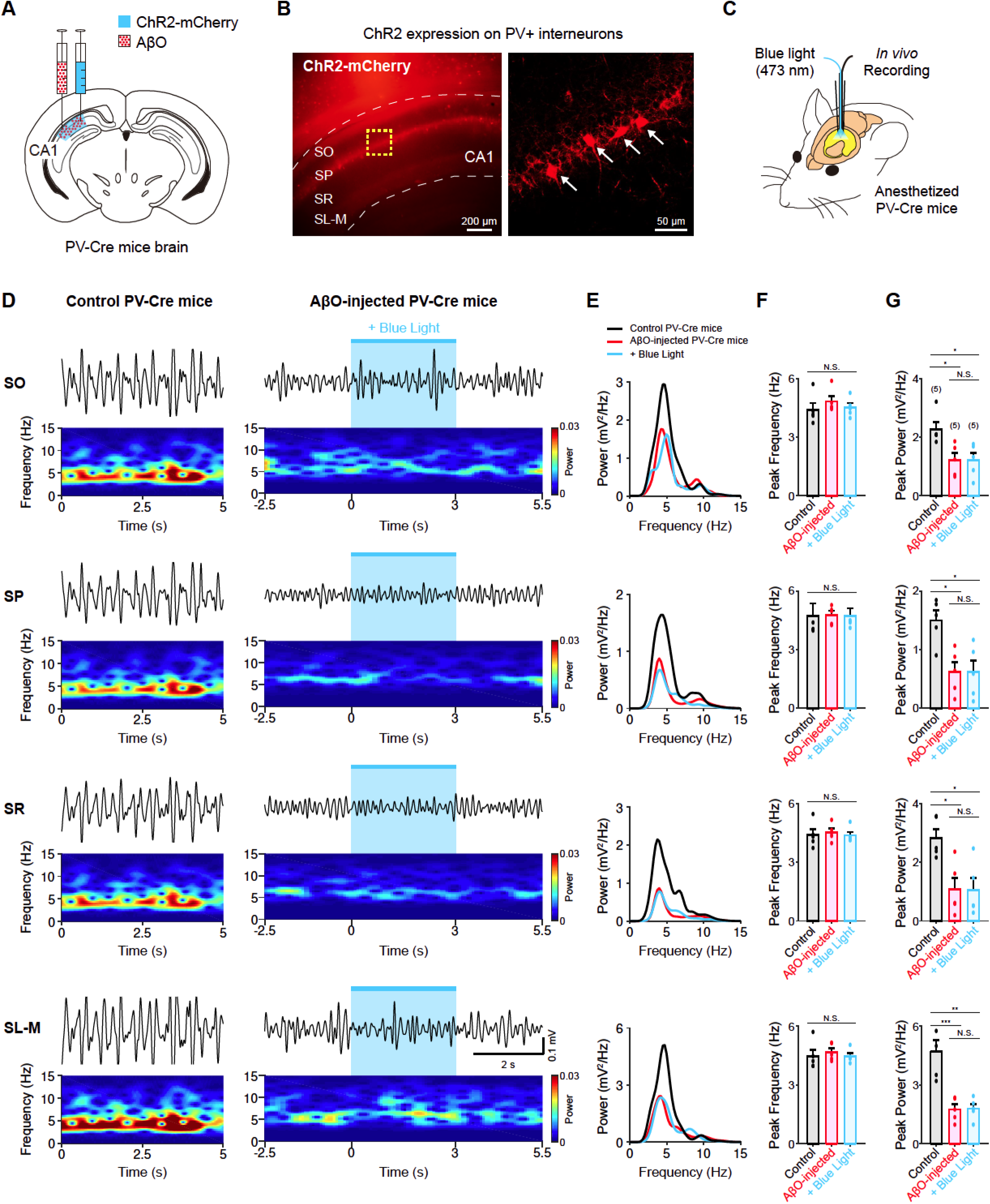
Optogenetic activation of PV+ interneurons had no effect on impaired theta oscillation of AβO-injected PV-Cre mice. (A) Schematic illustration of micro-injection of AβO and ChR2 virus into hippocampal CA1 region of PV-Cre mice. (B) Fluorescence image of ChR2 expressed PV+ interneurons in hippocampal CA1 region. Individual neurons are pointed by white arrows. (C) Schematic illustration of *in vivo* recording. (D) Example traces of spontaneous theta-filtered LFP signals (top) and spectrogram (bottom) recorded in control PV-Cre mice (left) and AβO-injected PV-Cre mice (right). The 3 s-long blue light stimulation of PV+ interneuron**s** is represented by light blue shaded region. (E) Example traces of PSD curves obtained from the spontaneous theta oscillation recorded in control PV-Cre mice (black), AβO-injected PV-Cre mice (red) and AβO-injected PV-Cre mice with blue light (blue). (F-G) Mean peak frequency (F) and mean peak power (G) of spontaneous theta-filtered LFP signals of control PV-Cre mice (black, n = 5), AβO-injected PV-Cre mice (red, n = 5) and AβO-injected PV-Cre mice with blue light (blue, n = 5). All data represent mean ± SEM. Inset: N.S. *p* > 0.05, ^∗^ *p* < 0.05, ^∗∗^ *p* < 0.01 and ^∗∗∗^ *p* < 0.001, one-way ANOVA followed by Tukey’s *post hoc* test.

### Hippocampal *in vivo* recording

Mice were deeply anesthetized with a mixture of ketamine (75-100 mg/kg) and medetomidine (1 mg/kg) and head-fixed into a stereotaxic frame (Stoelting Co.). *In vivo* local field potential (LFP) recordings were performed using a 32-channel silicon probe (A1×32-Poly2-5mm-50s-177, Neuronexus) inserted into the hippocampal CA1 region (2.7 mm posterior, 2.7 mm lateral to bregma, and 1.85 mm ventral to the pia). A reference metal wire was inserted into cerebellum. Body temperature was monitored and maintained at ~37 □ using a DC temperature control system (FHC Inc.). Spontaneous LFP signals and spontaneous spiking activities were recorded at a sampling rate of 1 kHz and 25 kHz (RZ2, Tucker-Davis Technologies).

For the optogenetic activation of SST+ or PV+ interneurons, 473 nm blue LED light stimulation (X-Cite 110LED, Excelitas Technologies, 50% light intensity) was delivered via an optical fiber laminated on the 32-channel silicon probe with the fiber tip terminating 100 μm above the top-most electrode. During a recording session, 473 nm blue light was delivered to CA1 area of the hippocampus for 3 s or 10 s at a time with an inter-trial interval of 60 s, repeated 10 times. The light stimulation protocol was controlled by custom-made pulse generator (Arduino).

### Immunochemistry

To verify that AβO was properly injected into the hippocampal CA1 region, immunochemistry was performed to detect AβO in hippocampal tissue. The 300 μm hippocampal slices were obtained from AβO-injected mice and fixed in 4% paraformaldehyde (Sigma-Aldrich) for > 24 hours at 4 □. Immunostaining was performed on free-floating sections. Fixed slices were rinsed in wash buffer (0.3 % Triton X-100 in 0.1 M PBS) three times (15 min / wash). To deactivate endogenous peroxidase, slices were incubated in peroxidase buffer (0.3% H_2_O_2_ in 0.1 M PBS) for 20 min and washed again in wash buffer three times (15 min / wash). Antigens which were not targeted were blocked by incubation in 6% bovine serum albumin (BSA) and 0.3% Triton X-100 in 0.1 M PBS for > 24 hours at 4 □ and subsequently incubated with the primary antibody (Anti-beta Amyloid 1-42 antibody [mOC64]-Conformation-Specific, Abcam, 1:300 dilution in 0.1 M PBS) at 4 □ for 4 days. Slices were washed and incubated with secondary antibody (Cy™3 AffiniPure Donkey Anti-Rabbit IgG (H+L), Jackson ImmunoResearch, 1:500 dilution in 0.1 M PBS). All slices were washed and mounted on slide glasses with cubic mount (150 g of 50% sucrose, 125 g of 25% urea, 125 g of 25% N, N, N’, N’-tetrakis(2-hydroxypropyle)ethlenediamine and 150 mL distilled water; (Lee et al., 2016)) and coverslipped. Immunostained AβO in hippocampal slices was visualized with a fluorescent microscope (Leica DM2500, Fig. 1B).

We confirmed the ChR2 expression in SST+ and PV+ interneurons by detecting the fluorescent signal of mCherry fluorophore under fluorescent microscope (Fig. 1C and Fig. 2B). To verify the position of the implanted probe in the hippocampus and identify the location of each electrode of the probe in each hippocampal layer, the electrode was removed after *in vivo* recording and coated with a fluorescent dye (Alexa 594, Fig. 1E), after which it was re-implanted at the same *in vivo* recording coordination for fluorescent imaging.

### Data Analysis of *in vivo* LFPs and spikes

Acquired spontaneous LFP signals were analyzed using customized protocols in MATLAB (R2018a). LFP signals were analyzed using a common average filter and 3^rd^ order butterworth bandpass filter (3 - 12 Hz, Fig. 1F and Fig. 2D), after which fast Fourier transform (FFT) was performed to analyze the power spectrum density (PSD, Fig. 1G and Fig. 2E). The peak frequency and peak power were taken to characterize theta oscillation (Fig. 1H-I and Fig. 2F-G). For each animal, 3 slong LFP epochs were detected for the analysis.

To investigate the phase relationship between CA1 pyramidal neurons and theta-frequency oscillation in each AβO–injected SST-Cre and PV-Cre mice, we first extracted single unit spikes by performing spike detection and sorting using the Klusta-suite software (Rossant et al., 2016). Spike events were detected when the 300-5,000 Hz band-pass filtered signal exceeds 4.5 times the standard deviation. Spike waveform, inter-spike intervals (ISIs), and the shape of auto-correlogram of spike times were examined to refine single unit identification (Hill et al., 2011). We classified putative CA1 pyramidal neurons and putative CA1 interneurons by calculating the asymmetry index ([(b - a)/(b + a)]) from the spike waveform (Fig. 3A). Neurons that were located on the right of the decision boundary ([(b - a)/(b + a)] = 2^∗^c – 1.2) were considered putative CA1 pyramidal neurons (Fig. 3A). To confirm successful optogenetic activation of SST+ or PV+ interneurons with blue light, we calculated the peri-stimulus time histogram (PSTH, 100 ms bin) before and after light stimulation (Fig. 3B). To investigate the spike phases of CA1 pyramidal neurons relative to theta oscillation, the spike timings of CA1 pyramidal neurons were extracted and the corresponding phase of the theta oscillation for each spike time was analyzed using a Hilbert transform (Khodagholy et al., 2017; Nakazono et al., 2017; Tort et al., 2010). The probability distribution of spike phases was normalized by dividing the bins (10 degree) by the total number of spikes (Fig. 3D, G). Each spike phase probability was vectorized on the polar coordinate. Average of spike phases was analyzed using the Circular Statistics Toolbox in MATLAB (Berens, 2009) (Fig. 3D, G). To quantify the strength of phase-locking, we obtained the vector of each spike phases of each CA1 pyramidal neurons and calculated the mean vector length (Fig. 3E, H).

**Fig. 3.**
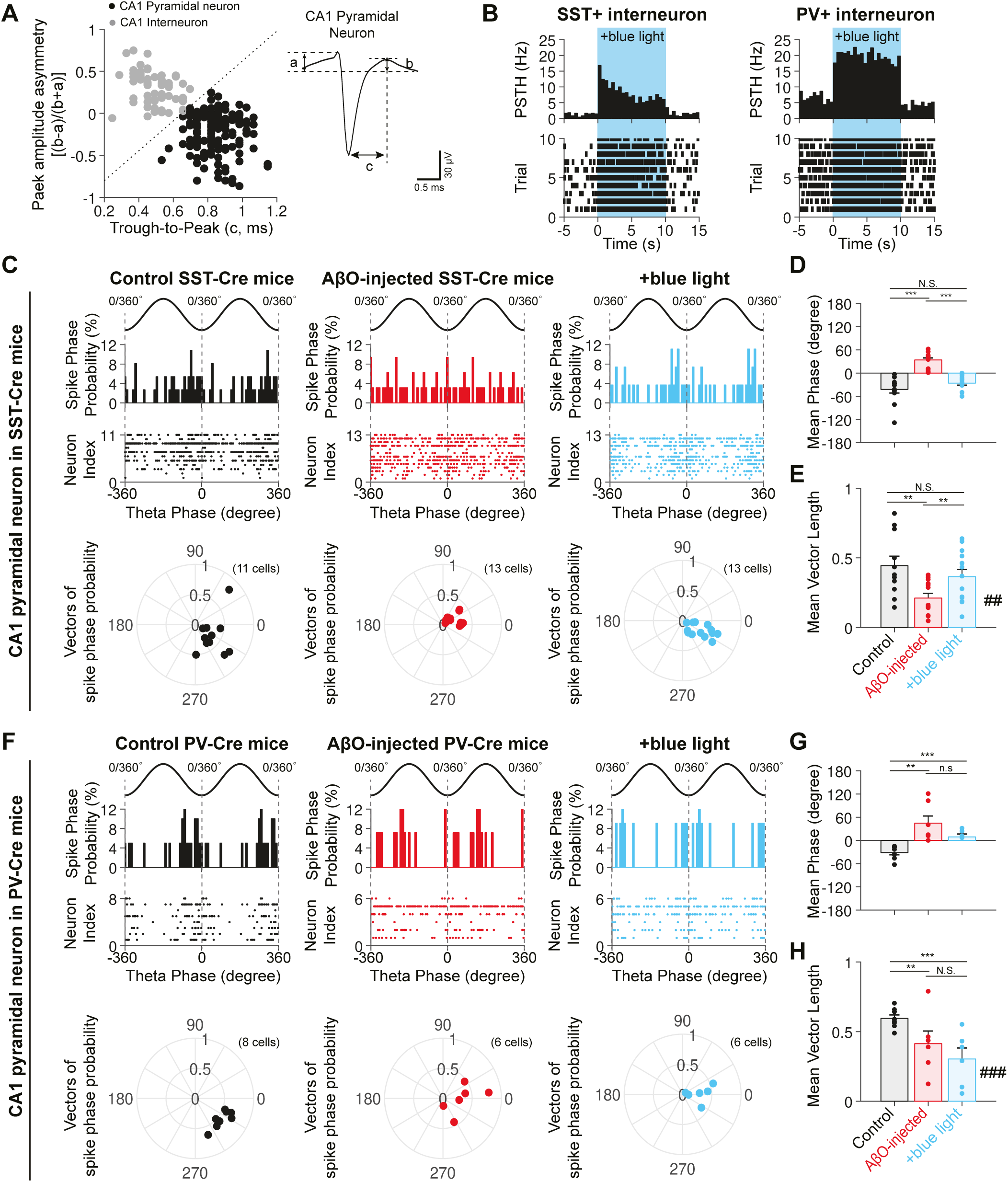
Optogenetic activation of SST+ interneurons restores phase-locking of CA1 pyramidal neurons relative to theta oscillation in AβO-injected PV-Cre mice. (A) Classification (left) of putative CA1 pyramidal neurons (black) and putative CA1 interneurons (gray) based on spike waveform asymmetry ([(b-a)/(b+a)]) to the trough-to-peak latency (c), and an example spike waveform of CA1 pyramidal neuron (right). (B) Peri-stimulus histogram (PSTH, top) and raster plot (bottom) of SST+ interneuron (left) and PV+ interneuron (right) during blue light stimulation (+blue light). (C) Probability distribution of spike time to theta phase (spike phase, top), raster plot (middle), and polar plot showing spike phases and vector lengths in control SST-Cre mice (black), AβO-injected SST-Cre mice (red), and AβO-injected SST-Cre mice with blue light (blue). (D-E) Circular average of spike phases (D) and mean vector length of spike phase probability (E) in control SST-Cre mice (black), AβO-injected SST-Cre mice (red), and AβO-injected SST-Cre mice with blue light (blue). (G-I) Same as (D-F) but in PV-Cre mice. All data represent mean ± SEM. Inset: N.S. *p* > 0.05, ^∗^, # *p* < 0.05, ^∗∗^, ## *p* < 0.01, ^∗∗∗^, ### *p* < 0.001, Watson-Williams multi-sample circular test for mean phases, one-way ANOVA followed by Tukey’s *post hoc* test for vector length.

### Brain slice preparation for *in vitro* recording

After three weeks of AβO and virus incubation following stereotaxic surgery, mice were anaesthetized using 1.25% Avertin solution (0.8 g of 2, 2, 2-Tribromoethanol, 0.5 ml of 2-methyl-2-butanol and 39.5 mL saline, Sigma Aldrich; (Papaioannou and Fox, 1993)) and perfused with ice-cold cutting solution containing (in mM): 180 sucrose, 2.5 KCl, 1.25 NaH_2_PO_4_, 25 NaHCO_3_, 11 glucose, 2 MgSO_4_, and 1 CaCl_2_ at pH 7.2-7.4 oxygenated with 95% O_2_ / 5% CO_2_, after which the brain was removed in oxygenated ice-cold cutting solution following decapitation. Coronal hippocampal slices (300 μm) were cut using a vibratome (VT 1000 □ S, Leica Microsystems) and allowed to recover for 20 min in a solution made up of a mixture of cutting solution and aCSF containing (in mM): 126 NaCl, 3 KCl, 1.25 NaH_2_PO_4_, 2 MgSO_4_, 2 CaCl_2_, 25 NaHCO_3_, and 10 glucose, pH 7.2–7.4) at a 1:1 ratio, which was oxygenated with 95% O_2_ / 5% CO_2_ gas. Slices were further incubated in oxygenated aCSF solution for at least 1 hour before recording at 30-32 °C.

### *In vitro* whole-cell patch-clamp recordings

Inhibitory postsynaptic currents (IPSCs) evoked by SST+ or PV+ interneurons were recorded from visually identified CA1 pyramidal neurons (differential interference contrast microscopy, BW51W, Olympus) in whole-cell voltage-clamp recordings (holding potential: +10 mV) using a borosilicate glass electrode (4-8 MΩ) filled with intracellular solution ((in mM): 115 Cesium methanesulfonate (CsMSF), 8 NaCl, 10 HEPES, 0.3 Na_3_-GTP, 4 Mg-ATP, 0.3 EGTA, 5 QX-314, and 10 BAPTA (pH 7.3 - 7.4 and 280 - 290 mOsm, Fig. 4A and 4D). Ten pulses of light (473 nm blue LED light, 5 ms, 15 mW) were delivered to hippocampal slices prepared from SST-Cre or PV-Cre mice to evoke SST+ interneuron-driven IPSCs (IPSC_SST_, Fig. 4B) or PV+ interneuron-driven IPSCs (IPSC_PV_, Fig. 4E) in CA1 pyramidal neurons. All light stimulation was delivered using a digital micromirror device (Polygon400, Mightex) and through the objective (40 X) of an BX51 microscope (Olympus). We used Igor Pro (Wavemetrics) to control the digital micromirror device and synchronize the light delivery with the electrophysiological recordings. All *in vitro* data recordings were acquired at 5 kHz using an ITC-18 AD board (HEKA) controlled by Igor Pro. Igor Pro was used for generating command signals. Short-term plasticity of IPSC_SST_ and IPSC_PV_ was analyzed by normalizing the IPSC amplitudes in the train (IPSC_n_) to the first IPSC (IPSC_1_) amplitude in the train (IPSC_n_/IPSC_1_) (Fig. 4C and 4F).

**Fig. 4.**
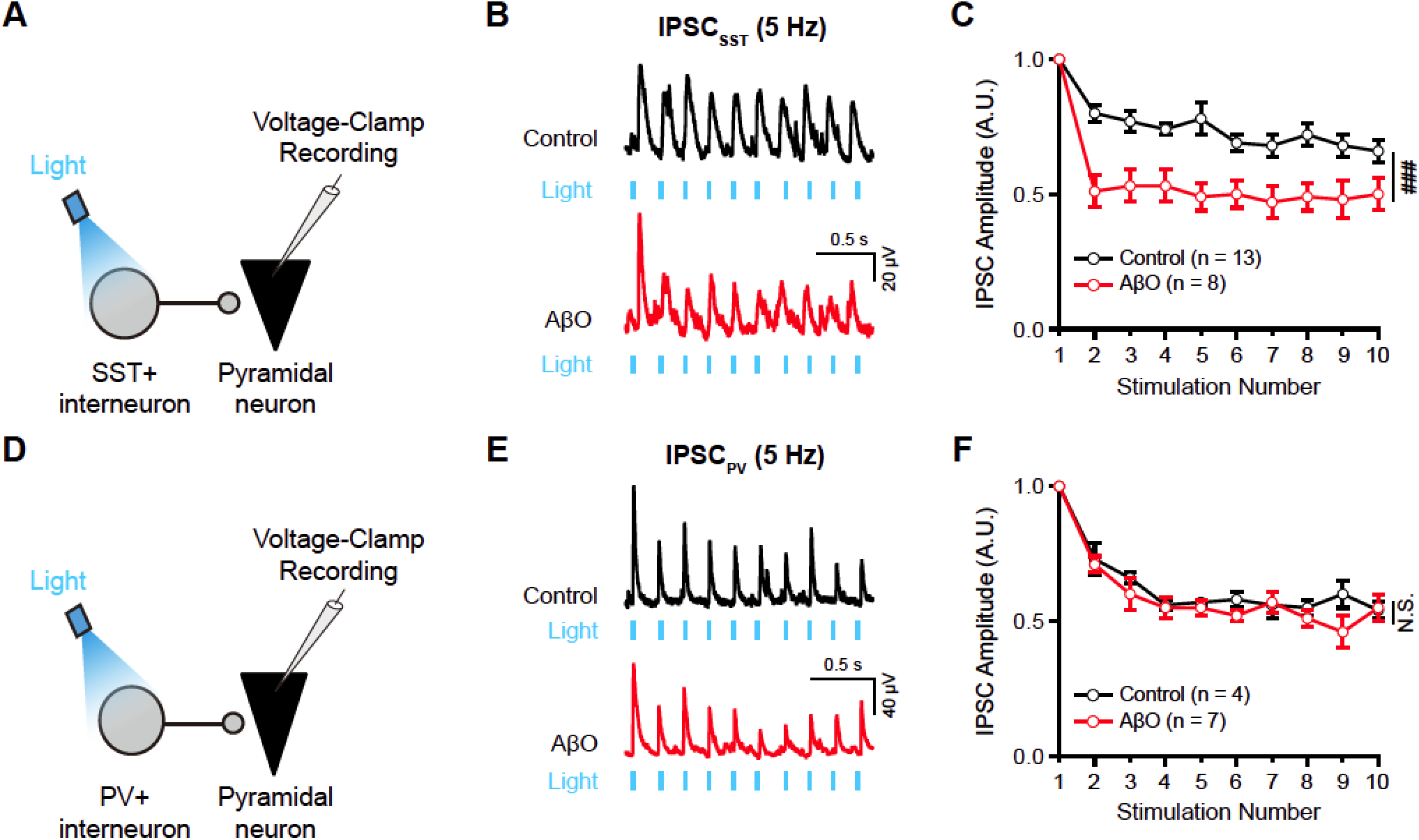
AβO causes selective synaptic dysfunction of SST+ interneuron inputs to CA1 pyramidal neuron at theta-frequency. (A) Schematic illustration of voltage-clamp recording of inhibitory postsynaptic current (IPSC) evoked by optical stimulation of SST+ interneuron (IPSC_SST_) in CA1 pyramidal neuron using blue light. (B) Example traces of IPSC_SST_ evoked by blue light stimulation at 5 Hz in DMSO-treated slice (control, top, black) and AβO-treated slice (AβO, bottom, red). (C) Short-term plasticity of IPSC_SST_ in control (black) and AβO slices (red, two-way ANOVA, control: n = 13, AβO: n = 8). (D) Schematic illustration of voltage-clamp recording of PV+ interneuron-driven IPSC (IPSC_PV_) onto CA1 pyramidal neuron using blue light stimulation. (E) Example traces of IPSC_PV_ by blue light stimulation at 5 Hz in the control (top, black) and AβO slice (bottom, red). (F) Short-term plasticity of IPSC_PV_ in control (black) and AβO slices (red, two-way ANOVA, control: n = 4, AβO: n = 7). All data represent mean ± SEM. Inset: ### *p* < 0.001, N.S. *p* > 0.05.

### Statistical analysis

All data are represented as mean ± standard error of the mean (SEM). Statistical analysis was performed by one-way and two-way ANOVA followed by Tukey’s *post hoc* test. *p* values less than 0.05 were considered statistically significant. Statistical significance of spike phases were tested using Watson-Williams multi-sample circular test (Zar, 2009).

## Results

In order to investigate the effect of optogenetic activation of SST+ interneurons on theta-frequency oscillation impairment observed in Alzheimer’s disease, we created an AD mouse model by injecting AβO into the CA1 area of the hippocampus of SST-Cre mice under anesthesia *in vivo* and co-injected ChR2 virus (AAV5-Ef1a-DIO-hChR2(ET/TC)-mCherry) (Fenno et al., 2011) to express ChR2 specifically in SST+ interneurons in mice with AβO pathology (Fig. 1A). We allowed three weeks for AβO and virus incubation following injections, after which we could clearly visualize deposition of AβO across all layers of the hippocampal CA1 area (Fig. 1B) and successful expression of ChR2 in SST+ interneurons in CA1 area of AβO-injected SST-Cre mice (Fig. 1C). To characterize the hippocampal theta-frequency oscillation in the control SST-Cre mice without AβO injection (control SST-Cre mice) and AβO-injected SST-Cre mice, we recorded spontaneous LFP under ketamine/medetomidine-induced anesthesia *in vivo* using an optrode, which is a 32-channel silicone probe laminated with an optical fiber that can simultaneously record LFP while delivering blue light (473 nm, Fig. 1D). The location of the electrode was confirmed by staining the recording location with Alexa 594 fluorescent dye after LFP recordings. Only recordings where the electrode track was visible in all four distinct layers of the CA1 area, stratum oriens (SO), stratum pyramidale (SP), stratum radiatum (SR) and stratum lacunosum-moleculare (SL-M), were used in further analysis (Fig. 1E). Band-pass filtered LFP in theta-frequency range (3 - 12 Hz, Fig. 1F top) and the corresponding spectrograms (Fig. 1F bottom) across all four layers in the control SST-Cre mice, AβO-injected SST-Cre mice and AβO-injected SST-Cre mice with blue light (473 nm) were directly compared as shown in Fig. 1F. We observed that the power of theta oscillations in control SST-Cre mice (Fig. 1F left) was reduced in AβO-injected SST-Cre mice (Fig. 1F right), which was increased to a level similar to that in the control mice during 3 s-long blue light stimulation of SST+ interneurons (Fig. 1F light blue shaded region). To quantify the changes in theta oscillations observed in Fig. 1F, we performed power spectral density (PSD) analysis (Fig. 1G) and analyzed the peak frequency and the peak power of theta oscillations. We found that the peak frequency of theta oscillations across all four layers remained stable in all three experimental conditions (SO: control SST-Cre mice = 4.16 ± 0.14 Hz (n = 5), AβO-injected SST-Cre mice = 4.58 ± 0.26 Hz (n = 9), AβO-injected SST-Cre mice + blue light = 4.69 ± 0.23 Hz (n = 9), F(2,20) = 1.02, *p* = 0.38; SP: control SST-Cre mice = 4.13 ± 0.21 Hz (n = 5), AβO-injected SST-Cre mice = 4.92 ± 0.28 Hz (n = 9), AβO-injected SST-Cre mice + blue light = 4.71 ± 0.20 Hz (n = 9), F(2,20) = 2.17, *p* = 0.14; SR: control SST-Cre mice = 4.40 ± 0.27 Hz (n = 5), AβO-injected SST-Cre mice = 4.82 ± 0.23 Hz (n = 9), AβO-injected SST-Cre mice + blue light = 5.09 ± 0.40 Hz (n = 9, F(2,20) = 0.92, *p* = 0.41; SL-M: control SST-Cre mice = 4.38 ± 0.19 Hz (n = 5), AβO-injected SST-Cre mice = 4.47 ± 0.22 Hz (n = 9), AβO-injected SST-Cre mice + blue light = 4.55 ± 0.27 Hz (n = 9), F(2,20) = 0.10, *p* = 0.90; one-way ANOVA; Fig 1H). However, peak power of theta oscillation in the control SST-Cre mice significantly decreased in all layers of AβO-injected SST-Cre mice, which was restored by optogenetic stimulation of SST+ interneurons in AβO-injected SST-Cre mice with blue light (SO: control SST-Cre mice = 1.14 ± 0.13 mV^2^/Hz (n = 5), AβO-injected SST-Cre mice = 0.48 ± 0.10 mV^2^/Hz (n = 9), AβO-injected SST-Cre mice + blue light = 0.98 ± 0.18 mV^2^/Hz (n = 9) (control *vs.* AβO-injected SST-Cre mice: ^∗^ *p* < 0.05, AβO-injected SST-Cre mice *vs.* AβO-injected SST-Cre mice + blue light: ^∗^ *p* < 0.05, control *vs.* AβO-injected SST-Cre mice + blue light: *p* = 0.76); SP: control SST-Cre mice = 0.83 ± 0.16 mV^2^/Hz (n = 5), AβO-injected SST-Cre mice = 0.33 ± 0.07 mV^2^/Hz (n = 9), AβO-injected SST-Cre mice + blue light = 0.72 ± 0.10 mV^2^/Hz (n = 9) (control *vs.* AβO-injected SST-Cre mice: ^∗^ *p* < 0.05, AβO-injected SST-Cre mice *vs.* AβO-injected SST-Cre mice + blue light: ^∗^ *p* < 0.05, control *vs.* AβO-injected SST-Cre mice + blue light: *p* = 0.77); SR: control SST-Cre mice = 1.08 ± 0.24 mV^2^/Hz (n = 5), AβO-injected SST-Cre mice = 0.36 ± 0.05 mV^2^/Hz (n = 9), AβO-injected SST-Cre mice + blue light = 0.84 ± 0.14 mV^2^/Hz (n = 9) (control *vs.* AβO-injected SST-Cre mice: ^∗∗^ *p* < 0.01, AβO-injected SST-Cre mice *vs.* AβO-injected SST-Cre mice + blue light: ^∗^ *p* < 0.05, control *vs.* AβO-injected SST-Cre mice + blue light: *p* = 0.49); SL-M: control SST-Cre mice = 3.22 ± 0.67 mV^2^/Hz (n = 5), AβO-injected SST-Cre mice = 1.31 ± 0.16 mV^2^/Hz (n = 9), AβO-injected SST-Cre mice + blue light = 2.75 ± 0.46 mV^2^/Hz (n = 9) (control *vs.* AβO-injected SST-Cre mice: ^∗^ *p* < 0.05, AβO-injected SST-Cre mice *vs.* AβO-injected SST-Cre mice + blue light: ^∗^ *p* < 0.05, control *vs.* AβO-injected SST-Cre mice + blue light: *p* = 0.74); one-way ANOVA followed by Tukey’s *post hoc* test; Fig. 1I). These results show that AβO-injection-induced decrease in peak power of hippocampal theta oscillations can be fully restored to control levels by selective optogenetic activation of SST+ interneurons in AβO-injected SST-Cre mice.

We next investigated the effect of another major interneuron subtype in the hippocampus, PV+ interneurons, on theta-frequency oscillation impairment observed in AD, by repeating the same method in Fig. 1 but now in PV-Cre mice (Fig. 2A-C). Band-pass filtered LFP in theta-frequency range (3 – 12 Hz, Fig. 2D top) and the corresponding spectrograms (Fig. 2D bottom) recorded across all four layers in the control PV-Cre mice, AβO-injected PV-Cre mice and AβO-injected PV-Cre mice with blue light (473 nm) were directly compared as shown in Fig. 2D. We observed that the power of theta oscillation in the control PV-Cre mice (Fig. 2D left) was reduced in AβO-injected PV-Cre mice (Fig. 2D right), which remained unchanged during 3 s-long blue light stimulation of PV+ interneurons with blue light (Fig. 2D right light blue shaded region). PSD analysis (Fig. 2E) revealed that the peak frequency of theta oscillations across all four layers remained stable in all three experimental conditions (SO: control PV-Cre mice = 4.40 ± 0.34 Hz (n = 5), AβO-injected PV-Cre mice = 4.82 ± 0.27 Hz (n = 5), AβO-injected PV-Cre mice + blue light = 4.52 ± 0.26 Hz (n = 5), F(2,12) = 0.60, *p* = 0.57; SP: control PV-Cre mice = 4.73 ± 0.62 Hz (n = 5), AβO-injected PV-Cre mice = 4.80 ± 0.19 Hz (n = 5), AβO-injected PV-Cre mice + blue light = 4.67 ± 0.39 Hz (n = 5), F(2,12) = 0.02, *p* = 0.98; SR: control PV-Cre mice = 4.40 ± 0.29 Hz (n = 5), AβO-injected PV-Cre mice = 4.53 ± 0.20 Hz (n = 5), AβO-injected PV-Cre mice + blue light = 4.37 ± 0.18 Hz (n = 5), F(2,12) = 0.13, *p* = 0.88; SL-M: control PV-Cre mice = 4.47 ± 0.32 Hz (n = 5), AβO-injected PV-Cre mice = 4.65 ± 0.22 Hz (n = 5), AβO-injected PV-Cre mice + blue light = 4.42 ± 0.16 Hz (n = 5), F(2,12) = 0.24, *p* = 0.79; one-way ANOVA; Fig 2F). Peak power of theta oscillation in the control PV-Cre mice significantly decreased in all layers of AβO-injected PV-Cre mice but, in contrast to optogenetic activation of SST+ interneurons, optogenetic stimulation of PV+ interneurons in AβO-injected PV-Cre mice had no effect on the peak power of theta oscillations, with the peak power remaining the same as before light stimulation (SO: control PV-Cre mice = 2.26 ± 0.24 mV^2^/Hz (n = 5), AβO-injected PV-Cre mice = 1.22 ± 0.25 mV^2^/Hz (n = 5), AβO-injected PV-Cre mice + blue light = 1.21 ± 0.25 mV^2^/Hz (n = 5) (control *vs.* AβO-injected PV-Cre mice: ^∗^ *p* < 0.05, AβO-injected PV-Cre mice *vs.* AβO-injected PV-Cre mice + blue light: *p* = 1.00, control *vs.* AβO-injected PV-Cre mice + blue light: ^∗^ *p* < 0.05); SP: control PV-Cre mice = 1.50 ± 0.17 mV^2^/Hz (n = 5), AβO-injected PV-Cre mice = 0.62 ± 0.17 mV^2^/Hz (n = 5), AβO-injected PV-Cre mice + blue light 0.62 ± 0.20 mV^2^/Hz (n = 5) (control *vs.* AβO-injected PV-Cre mice: ^∗^ *p* < 0.05, AβO-injected PV-Cre mice *vs.* AβO-injected PV-Cre mice + blue light: *p* = 1.00, control *vs.* AβO-injected PV-Cre mice + blue light: ^∗^ *p* < 0.05); SR: control = 2.83 ± 0.30 mV^2^/Hz (n = 5), AβO-injected PV-Cre mice = 1.07 ± 0.40 mV^2^/Hz (n = 5), AβO-injected PV-Cre mice + blue light = 1.05 ± 0.41 mV^2^/Hz (n = 5) (control *vs.* AβO-injected PV-Cre mice: ^∗^ *p* < 0.05, AβO-injected PV-Cre mice *vs.* AβO-injected PV-Cre mice + blue light: *p* = 0.99, control *vs.* AβO-injected PV-Cre mice + blue light: ^∗^ *p* < 0.05); SL-M: control PV-Cre mice = 4.70 ± 0.58 mV^2^/Hz (n = 5), AβO-injected PV-Cre mice = 1.74 ± 0.27 mV^2^/Hz (n = 5), AβO-injected PV-Cre mice + blue light = 1.78 ± 0.24 mV^2^/Hz (n = 5) (control vs. AβO-injected PV-Cre mice: ^∗∗∗^ *p* < 0.001, AβO-injected PV-Cre mice *vs.* AβO-injected PV-Cre mice + blue light: *p* = 0.99, control vs. AβO-injected PV-Cre mice + blue light: ^∗∗^ *p* < 0.01); one-way ANOVA followed by Tukey’s *post hoc* test; Fig. 2G). These results show that optogenetic stimulation of PV+ interneurons during hippocampal theta oscillation in AβO-injected PV-Cre mice had no detectable effect on theta oscillation, further underscoring the involvement of SST+ interneurons in theta oscillogenesis.

Hippocampal theta oscillation is used as reference for hippocampal spike phase codes where CA1 pyramidal neurons spike at specific phases of theta oscillation cycle, thought to be critical for spatial navigation and memory functions (Dragoi and Buzsaki, 2006; Maurer and McNaughton, 2007; O’Keefe and Recce, 1993; Skaggs et al., 1996). Therefore, we investigated the effect of optogenetic activation of SST+ and PV+ interneurons on theta spike phases in AβO-injected SST-Cre and PV-Cre mice, respectively. Firstly, we isolated single spikes of neurons from the band-pass (300-5,000 Hz) filtered signal and divided spikes into putative CA1 pyramidal neurons and CA1 interneurons based on the spike waveforms (Fig 1A), spike inter-spike intervals (ISIs) and autocorrelogram (see Methods). We confirmed successful optogenetic activation of SST+ and PV+ interneurons in SST-Cre and PV-Cre mice as illustrated by the increase in spike firing rates in putative SST+ interneurons and PV+ interneurons in the peri-stimulus time histogram (PSTH, Fig. 3B). Only spikes from putative CA1 pyramidal neurons were used in the analysis of theta-spike phases. To quantify the strength of synchronization of CA1 pyramidal neurons’ spikes, we first analyzed the probability distribution of spike phases of CA1 pyramidal neurons relative to theta cycle (Fig. 3C top), which were represented in the polar plot (Fig. 3C bottom) in the control SST-Cre mice (Fig. 3C, black), AβO-injected SST-Cre mice (Fig. 3C, red) and AβO-injected SST-Cre mice with blue light (Fig. 3C blue). Analysis of mean spike phases relative to the theta cycle revealed that CA1 pyramidal neurons spiked preferentially before the trough of theta cycle (−42.14 ± 9.47 degrees, n = 11 cells; Fig. 3D, black), consistent with previous *in vivo* recordings in anesthetized mice (Huh et al., 2016; Klausberger et al., 2003; Somogyi et al., 2014). However, theta spike phases of CA1 pyramidal neurons in AβO-injected SST-Cre mice occurred at a significantly later phase after the trough of the oscillation compared to the control case (33.89 ± 5.38 degrees, n = 13 cells; Fig. 3D, red). Optical stimulation of SST+ interneurons in AβO-injected SST-Cre mice restored the spike phases similar to that in the control SST-Cre mice (−25.98 ± 5.29 degrees, n = 13 cells, Fig 3D blue; control SST-Cre mice *vs.* AβO-injected SST-Cre mice: ^∗∗∗^ *p* < 0.001; AβO-injected SST-Cre mice *vs.* AβO-injected SST-Cre mice + blue light: ^∗∗∗^ *p* < 0.001; control SST-Cre mice *vs.* AβO-injected SST-Cre mice + blue light: N.S. *p* > 0.05, Watson-Williams multi-sample circular test). We also found that the mean vector length of probability distribution of spike phases was significantly decreased in AβO-injected SST-Cre mice compared to the control SST-Cre mice suggesting that CA1 pyramidal neurons’ spikes were significantly desynchronized in AβO-injected SST-Cre mice and light stimulation of SST+ interneurons in AβO-injected SST-Cre mice fully restored the vector length to the level of that observed in the control SST-Cre mice (control SST-Cre mice = 0.44 ± 0.07 (n = 11 cells), AβO-injected SST-Cre = 0.21 ± 0.03 (n = 13 cells), AβO-injected SST-Cre + blue light = 0.37 ± 0.05 (n = 13 cells); ## *p* < 0.01; control SST-Cre mice vs. AβO-injected SST-Cre mice: ^∗∗^ *p* < 0.01; AβO-injected SST-Cre mice vs. AβO-injected SST-Cre mice + blue light: ^∗∗^ *p* < 0.01; control SST-Cre mice vs. AβO-injected SST-Cre mice + blue light: N.S. *p* > 0.05; one-way ANOVA followed by Tukey’s *post*-*hoc* test; Fig. 3E). When we repeated the same analysis now on control PV-Cre mice, AβO-injected PV-Cre mice, and AβO-injected PV-Cre mice stimulated with blue light (Fig. 3F), we found no detectable changes in neither mean spike phases (control PV-Cre mice = −31.43 ± 5.55 degrees (n = 8 cells), AβO-injected PV-Cre mice = 44.83 ± 18.20 degrees (n = 6 cells), AβO-injected PV-Cre mice + blue light = 13.78 ± 4.27 degrees (n = 6 cells); control PV-Cre mice *vs.* AβO-injected PV-Cre mice: ^∗∗^ *p* < 0.01; AβO-injected PV-Cre mice vs. AβO-injected PV-Cre mice + blue light: N.S. *p* > 0.05; control *vs.* AβO-injected + blue light PV-Cre: ^∗∗∗^ *p* < 0.001; Watson-Williams multi-sample circular test; Fig. 3G) nor vector length (control PV-Cre mice = 0.60 ± 0.02 (n = 8 cells), AβO-injected PV-Cre mice = 0.29 ± 0.07 (n = 6 cells), AβO-injected PV-Cre mice + blue light = 0.27 ± 0.08 (n = 6 cells); ### *p* < 0.001, one-way ANOVA; control vs. AβO-injected PV-Cre: ^∗∗^ *p* < 0.01; AβO-injected PV-Cre mice vs. AβO-injected PV-Cre mice + blue light: N.S. *p* > 0.05; control PV-Cre mice vs. AβO-injected PV-Cre mice + blue light: ^∗∗^ *p* < 0.01; one-way ANOVA followed by Tukey’s *post*-*hoc* test; Fig. 3H). These results show that optogenetic activation of SST+ interneurons in AβO-injected SST-Cre mice aids in resynchronizing the CA1 pyramidal neurons’ theta spike phases.

What could be the cellular mechanism underlying this restoration of synchrony? Since light stimulation of SST+ interneurons could selectively restore theta oscillation as well as the synchronization of theta spike phases in AβO-injected SST-Cre mice (Fig. 3), we hypothesized that light stimulation of SST+ interneuron restored dysfunctional SST+ interneuron inputs to CA1 pyramidal neurons in AβO-injected SST-Cre mice. In order to test this, we performed whole-cell voltage-clamp recordings on CA1 pyramidal neurons in hippocampal slices *in vitro* to measure inhibitory postsynaptic currents (IPSCs) in CA1 pyramidal neurons. IPSCs were evoked by blue light stimulation of ChR2-expressing SST (IPSC_SST_) in the hippocampal slices cut from SST-Cre mice and with either treated with DMSO (control) or AβO (Fig. 4A). When SST+ interneurons were stimulated with blue light at 5 Hz for 10 times in the control slices, IPSC_SST_ amplitude decreased as the stimulation pulse number increased, showing robust short-term depression (STD, Fig. 4B, black) while the same stimulation showed significant enhancement of STD in AβO-treated slices (Fig. 4B, red; Control (n = 13) *vs.* AβO (n = 8), ### *p* < 0.001, two-way ANOVA; Fig. 4C). In contrast, when the same experiment was repeated in PV-Cre mice to record IPSC in CA1 pyramidal neurons evoked by light stimulation of ChR2-expressing PV (IPSC_PV_) in hippocampal slices cut from PV-Cre mice treated with DMSO (control) and that treated with AβO (Fig. 4D), IPSC_PV_ amplitude decreased with increasing stimulation number showing robust STD in the control slices (Fig. 4E, black), which was unaffected in AβO-treated slices (Fig. 4E, red) as shown in statistical analysis (Control (n = 4) *vs.* AβO (n = 7), N.S. *p* > 0.05, two-way ANOVA; Fig. 4F). These results demonstrate for the first time that enhancement of STD in SST+ interneuronal input to CA1 pyramidal neurons in AβO-injected mice may underlie the desynchronization of CA1 pyramidal neuron output, leading to a reduction in theta oscillation power *in vivo.*

## Discussion

Here we show that optogenetic-activation of SST+ interneurons can selectively restore AβO-induced impairment of hippocampal theta oscillations and aid in resynchronizing theta spike phases of CA1 pyramidal neurons in mice. Such interneuron subtype-specific restoration of theta oscillation was possible since AβO caused synaptic dysfunction specifically to SST+ interneuron’s theta-frequency inhibitory inputs to CA1 pyramidal neurons, identifying SST+ interneurons as a target for restoring theta-frequency oscillations in early AD.

In our study, we injected AβO into the hippocampus to create AβO pathology *in vivo*, a method adopted in many studies investigating the impact of AβO physiological and cognitive functions (Balducci et al., 2010; Brouillette et al., 2012; Cetin and Dincer, 2007; Faucher et al., 2015; Kalweit et al., 2015; Kim et al., 2014b; Nicole et al., 2016; Orban et al., 2010; Villette et al., 2010; Yi et al., 2018). After three weeks of recovery following AβO injection in SST-Cre and PV-Cre mice, we observed a significant reduction in peak power of theta oscillations across all hippocampal sublayers compared to that in control SST-Cre and PV-Cre mice (Fig. 1-2). These results are consistent with the reduction of theta oscillation peak power observed in AβO-injected mice after three weeks of incubation (Villette et al., 2010).

The major advantage of using the AβO-injection model of AD in mice was that we were able to combined this with optogenetic activation of SST+ (Fig. 1F) and PV+ interneurons (Fig. 2D) using cre-transgenic mice. Using these mice, we were able to show, for the first time, that optogenetic activation of SST+ interneurons in AβO-injected mice could restore the power of theta oscillations (Fig. 1F-I) and CA1 pyramidal neuron’s spike output relative to theta oscillation phase (Fig. 3C-E) while optogenetic-activation of PV+ interneurons had no effect (Fig. 2D-G, 3F-H). Restoration of gamma oscillations in AD mice models has been demonstrated by modulating PV+ interneurons including optogenetic activation of PV+ interneuron (Iaccarino et al., 2016), implantation of PV-like fast-spiking interneurons (Tong et al., 2014) and even ablation of certain genes (Martinez-Losa et al., 2018; Verret et al., 2012; Zhang et al., 2017). However, to our knowledge, no study has yet demonstrated ways to restore theta oscillations in AD mouse models nor investigated the effect of optogenetic activation SST+ interneuron on hippocampal oscillations in AD.

We found that the spike phase of CA1 pyramidal neurons relative to theta oscillations, which normally occurs before the trough of the theta cycle, was significantly shifted to a later phase after the trough (Fig. 3C-D, 3F-G) and their strength of phase synchronization (or phase-locking) was significantly reduced in both AβO-injected SST-Cre and PV-Cre mice (Fig. 3E, H). These results are consistent with the same analysis performed in 3xTg AD mice models (Mably et al., 2017). However, we found that optogenetic activation of SST+ interneurons could specifically restore the desynchronized spike phase in AβO-injected SST-Cre mice (Fig. 3C-E) while optogenetic activation of PV+ interneurons had no effect on either of them (Fig. 3F-H).

Through *in vitro* whole-cell voltage-clamp recordings in CA1 pyramidal neurons in AβO-treated hippocampal slices, we found that STD of IPSCs, evoked by optically stimulating SST+ interneurons (IPSC_SST_) at theta frequency, was enhanced compared to that in control slices (Fig. 4A-C) while that evoked by optically stimulating PV (IPSC_PV_) was unaffected (Fig. 4D-F). These results indicate that the reduction in IPSC_SST_ to CA1 pyramidal neurons could explain desynchronized spikes, leading to the reduction of theta oscillation power and in AβO-injected mice.

Reduction in theta oscillation power has also been observed in other genetically modified AD mice models such as 3xTg (Mondragon-Rodriguez et al., 2018), CRND8 (Goutagny et al., 2013), and APP23 transgenic AD mice models (Ittner et al., 2014). However, in addition to the reduction of theta oscillation power, hypersynchrony and epileptic activities were also observed in transgenic AD mice models (Bezzina et al., 2015; Ittner et al., 2014; Palop and Mucke, 2010) which was absent in the AβO-injected mice in ours and other studies (Kalweit et al., 2015). Hence, long-term effects of AβO may cause differing effects on the hippocampal neural circuits, which will require further investigation. Also, since we performed all our *in vivo* recordings in animals under anesthesia i*n vivo*, it will be interesting to investigate how theta oscillations in the awake animal are affected by optogenetic activation of SST+ and PV+ interneurons in AβO-injected mice.

Overall, our results demonstrate for the first time that theta oscillation impairment through AβO pathology could arise from dysfunction of SST+ interneuron inputs to CA1 pyramidal neurons, causing desynchronization of CA1 pyramidal neuron’s spikes relative to theta oscillations, eventually leading to a reduction in theta oscillation power. These results provide experimental evidence how SST+ interneurons contribute to the theta oscillogenesis in the hippocampus and suggest that optogenetic modulation of SST+ interneurons could help restore hippocampal theta oscillation in early stages of AD.

## Acknowledgement

This work was supported by a grant of the Korea Health Technology R&D Project through the Korea Health Industry Development Institute (KHIDI), funded by the Ministry of Health & Welfare, Republic of Korea (HI15C3086, HI17C0212).

